# Tests for segregation distortion in higher ploidy F1 populations

**DOI:** 10.1101/2025.06.23.661114

**Authors:** David Gerard, Guilherme Bovi Ambrosano, Guilherme da Silva Pereira, Antonio Augusto Franco Garcia

## Abstract

F1 populations are widely used in genetic mapping studies in agriculture, where known pedigrees enable rigorous quality control measures such as segregation distortion testing. However, conventional tests for segregation distortion are inadequate for polyploids, as they fail to account for double reduction, preferential pairing, and genotype uncertainty, leading to inflated type I error rates. Gerard et al. (2025) (doi:10.1007/s00122-025-04816-z) developed a statistical framework to address these issues in tetraploids. Here, we extend these methods to higher even ploidy levels and introduce additional strategies to mitigate the influence of outliers. Through extensive simulations, we demonstrate that our tests maintain appropriate type I error control while retaining power to detect true segregation distortion. We further validate our approach using empirical data from a hexaploid mapping population. Our methods are implemented in the segtest R package, available on the GitHub (https://github.com/dcgerard/segtest).

## 1 Introduction

Experimental polyploid populations are frequently used in agricultural research for tasks such as linkage mapping, quantitative trait locus mapping, and genomic selection [Shirasawa et al., 2017, Bourke et al., 2018, Lara et al., 2019, Mollinari and Garcia, 2019, Pereira et al., 2020, Gemenet et al., 2020, Mollinari et al., 2020, Amadeu et al., 2021, Ferrão et al., 2021, Lau et al., 2022]. These sophisticated statistical methods require high-quality genotype data to achieve optimal performance. Consequently, genotype data are routinely subjected to rigorous quality control procedures [e.g., Bourke et al., 2015, Cappai et al., 2020, Mollinari et al., 2020, Batista et al., 2021]. Given that experimental populations possess well-defined family pedigrees, a frequent check involves using a *χ*^2^-test to compare observed offspring genotype frequencies with those predicted by Mendelian segregation [Mendel, 1866], as is done by the mappoly software [Mollinari et al., 2020]. However, this method has notable limitations: it fails to incorporate features specific to polyploid meiosis, such as double reduction and (partial) preferential pairing [Voorrips and Maliepaard, 2012], and does not account for the increased genotype uncertainty associated with modern sequencing technologies [Gerard et al., 2018, Gerard and Ferrão, 2019].

Recent approaches by Bourke et al. [2018] and Gerard et al. [2025] have been specifically tailored to polyploid data. However, both methods exhibit certain limitations. Bourke et al. [2018], through the checkF1() function of their polymapR software, implement a *χ*^2^-test for all polysomic and disomic segregation patterns in polyploids. Additionally, their method permits a specified proportion of individuals to have “invalid” genotypes (e.g., offspring genotypes greater than or equal to 2 are deemed invalid when the parent genotypes are 0 and 1). Despite these features, the approach of Bourke et al. [2018] has several shortcomings: (i) it does not account for partial preferential pairing or double reduction, (ii) it addresses genotype uncertainty in an ad hoc manner, which results in elevated type I error rates [Gerard et al., 2025], and (iii) it conducts separate tests for invalid genotypes and segregation distortion, an approach difficult to generalize when accounting for genotype uncertainty. See Gerard et al. [2025] for a detailed description of the checkF1() function.

Gerard et al. [2025] addressed some of these limitations by developing tests that incorporate both double reduction and partial preferential pairing, while systematically accounting for genotype uncertainty through the use of genotype likelihoods. However, the methods proposed by Gerard et al. [2025] are restricted to tetraploids and do not accommodate a proportion of invalid genotypes. Allowing for a certain degree of “wiggle room” in invalid genotypes can be important, as a SNP may be otherwise valid except for a few individuals with anomalous results. Discarding an entire SNP based solely on the presence of a few outliers could result in unnecessary data loss.

In this paper, we extend the work of Gerard et al. [2025] to develop tests for segregation distortion applicable to higher (even) ploidies, accounting for partial preferential pairing, invalid genotypes, and genotype uncertainty. Our approach integrates invalid genotypes into a single test for segregation distortion using a mixture model of genotype frequencies. Genotype uncertainty is addressed in a principled manner through the implementation of likelihood ratio tests (LRTs), utilizing genotype likelihoods. Our methods further account for the effects of double reduction through a novel result for their gamete frequencies at simplex loci, and we demonstrate that our methods are robust to moderate levels of double reduction at non-simplex loci.

## 2 Materials and methods

### 2.1 Models for genotype frequencies

In this section, we develop a null model for genotype frequencies in an F1 population, defining segregation distortion as deviations from these expected frequencies. The model includes a parameter *β* to account for double reduction at simplex loci, a parameter ***γ*** to account for partial preferential pairing (which is only an important modelling consideration at non-simplex loci), and a parameter *π* that specifies the tolerated proportion of outliers (i.e., “invalid genotypes”).

At a biallelic locus with alleles A and a, a *K*-ploid individual’s genotype is their number of a alleles. The genotype frequencies ***q*** = (*q*_0_*, q*_1_*, …, q_K_*) of an F1 population, where *q_k_* is the probability an offspring has genotype *k*, are given by the discrete linear convolution of parental gamete frequencies ***p****_j_* = (*p_j_*_0_*, p_j_*_1_*, …, p_j,K/_*_2_), where *p_jk_* is the probability parent *j*’s gamete has genotype *k*:

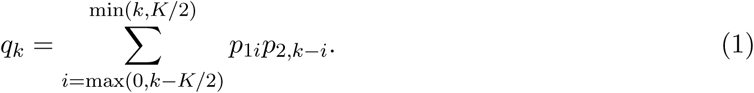

Different models for meiosis correspond to different models for the *p_jk_*’s.

For non-simplex loci, we account for (partial) preferential pairing using the pairing configuration model of Gerard et al. [2018]. During meiosis, a parent’s *K* chromosome copies form *K/*2 pairs. Let ***m*** = (*m*_0_*, m*_1_*, m*_2_) be the pairing configuration, where *m_j_*is the number of pairs containing *j* copies of a. Given a pairing configuration, the gamete frequencies are:

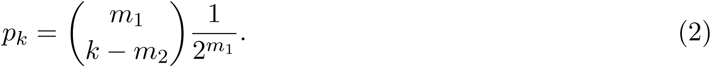

All possible values for the *p_k_*’s from (2) for ploidies 2 through 12 are presented in Table S1. Since the pairing configuration is unknown, let *γ_i_* denote the probability of pairing configuration *i* ∈ {1*, …, b_Kℓ_*}, where *b_Kℓ_* is the number of possible configurations for a parent with ploidy *K* and genotype *ℓ* [Gerard et al., 2018]:

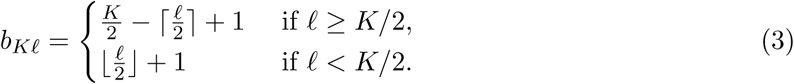

Marginalizing over pairing configurations, the gamete frequencies become:

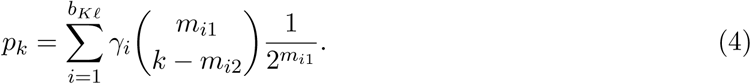

Model (4) generalizes the gamete frequencies of allopolyploids (disomic inheritance) and bivalent pairing autopolyploids (polysomic inheritance) [Doyle and Egan, 2010, Parisod et al., 2010]. For allopolyploids, ***γ*** is a one-hot vector (only one pairing configuration is possible). The value of ***γ*** that corresponds to bivalent pairing autopolyploids is given by Gerard et al. [2018]. For example, for autotetraploids where *ℓ* = 2, the probability of configuration ***m***_1_ = (1, 0, 1) is 1/3 and the probability of configuration ***m***_2_ = (0, 2, 0) is 2/3, resulting in an autotetraploid segregation pattern of

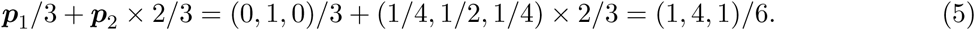

More generally, this model accommodates segmental allopolyploids with arbitrary levels of partial preferential pairing.

At simplex loci, only one pairing configuration exists (and so ***γ*** = 1). This simplifies modeling of double reduction at these loci. Theorem 1 defines genotype frequencies at simplex loci under arbitrary levels of double reduction (proof in Appendix S1). Upper bounds under two models for meiosis [Huang et al., 2019, Gerard, 2022] appear in Table S2.

**Theorem 1.** Let α_i_be the probability that there are i pairs of alleles in a gamete that are identical by double reduction. Then the gamete frequencies at a simplex locus are

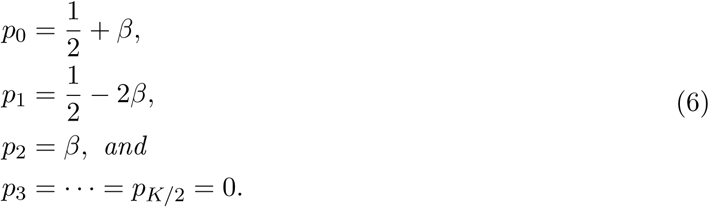

where

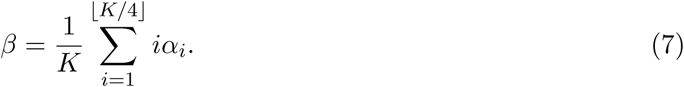

To incorporate invalid genotypes, we mix the genotype frequencies with a discrete uniform distribution:

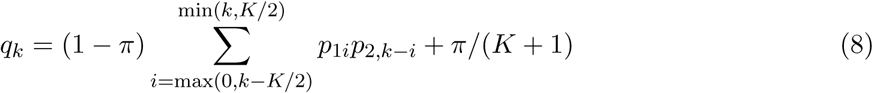

where *π* represents the proportion of offspring with anomalous genotypes. By default, we set *π* ≤ 0.03, but this is user-adjustable. This enables integrated modeling of genotype frequencies and outliers, unlike the two-step approach in polymapR [Bourke et al., 2018]. This makes it possible to model outliers in the presence of genotype uncertainty in Section 2.2.

In summary, we model the genotype frequencies as a mixture of the discrete uniform distribution and the convolution of gamete frequencies (8). The gamete frequencies can be specified by the user to correspond to full autopolyploidy, full allopolyploidy, or segmental allopolyploidy through different constraints on ***γ*** in (4). This model offers several advantages over polymapR [Bourke et al., 2018], including support for segmental allopolyploidy (which polymapR does not accommodate) and the direct incorporation of invalid genotypes into the model, rather than addressing them through a two-step procedure. Additionally, this approach extends the methods of Gerard et al. [2025] to higher ploidy levels while incorporating the ability to handle invalid genotypes. We account for double reduction at simplex markers, where the effects of partial preferential pairing are absent.

For clarity, in Appendix S2 we explicitly write out our model for the gamete frequencies for even ploidies 2 through 12.

### 2.2 Likelihood ratio tests for segregation distortion

In this section, we construct LRTs under the null hypothesis that the genotype frequencies correspond to one of the models of Section 2.1. We will construct these tests either using known genotypes or by accounting for genotype uncertainty through genotype likelihoods.

The likelihood for the model depends on whether genotypes are known or unknown. In the known genotype case, we let *x_k_* be the count of individuals with genotype *k* ∈ {0, 1*, …, K*}, which we organize into the vector ***x*** = (*x*_0_*, x*_1_*, …, x_K_*), with total sample size 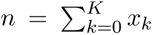. Given genotype frequencies ***q***, ***x*** follows a multinomial distribution, with the following likelihood

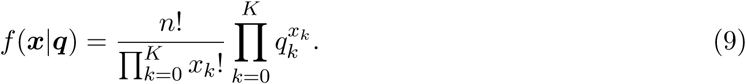

In the unknown genotype case, we represent genotype uncertainty through genotype likelihoods [Li, 2011]. Let *g_ik_* denote the genotype likelihood for individual *i* = 1, 2*, …, n* and genotype *k* = 0, 1*, …, K*. Specifically, *g_ik_* is the probability of the observed data (e.g., sequencing or microarray) for individual *i*, assuming their genotype is *k*. Given these genotype likelihoods, the likelihood function is

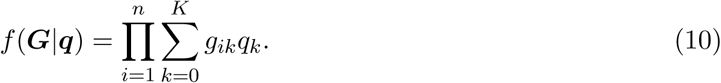

In the known genotype case (9), the maximum likelihood estimate of ***q*** under the alternative hypothesis is ***q***^*_A_* = ***x****/n*. In the unknown genotype case (10), ***q***^*_A_* can be calculated via the expectation-maximization (EM) algorithm of Li [2011]. By maximizing either (9) or (10) over the null parameter space, we obtain ***q***^_0_. Using this, we calculate the likelihood ratio statistic (2 times the difference of the maximized log-likelihoods under the null and alternative) and we compare it to an appropriate *χ*^2^ distribution to compute a *p*-value.

Our strategy to maximize over the null parameter space depends on the null hypothesis under consideration. If the null hypothesis corresponds to a true autopolyploid, the only unknown parameter is the invalid mixing proportion *π* in (8), which we maximize over using Brent’s method [Brent, 2013]. If the null hypothesis corresponds to a true allopolyploid (2), the two unknowns are the latent pairing configuration and the invalid mixing proportion *π* in (8). We iterate over each of the *b_Kℓ_*possible pairing configurations, optimizing over *π* via Brent’s method for each configuration, to identify the pairing configuration and *π* that maximize the likelihood. If the null hypothesis corresponds to a segmental allopolyploid (4), the unknowns include the mixing proportions for pairing configurations in both parents, ***γ***_1_ and ***γ***_2_, as well as the invalid genotype mixing proportion *π*. This optimization is performed using the method of Powell [2009], with initial values obtained via the global optimization method of Kucherenko and Sytsko [2005]. The simplex constraints on the mixing proportions for pairing configurations are incorporated using the parameterization of Betancourt [2012].

The number of degrees of freedom for the null *χ*^2^ distribution is the difference between the number of parameters under the alternative and under the null. The number of parameters under the alternative is just the ploidy of the offspring. Though, we can improve the power of the LRT if we subtract off the number of dosages whose frequencies are estimated to be 0 both under the null and under the alternative [Gerard et al., 2025].

Calculating the number of parameters under the null is tricky for two reasons: (i) some null parameters may lie on or near the boundary of the null parameter space, requiring extra consideration [Self and Liang, 1987, Mitchell et al., 2019, Leung and Sturma, 2024], and (ii) the null parameter space is weakly identified. To address the boundary problem (i), we adapt the data-dependent degrees of freedom strategy of Susko [2013]. Specifically, the number of parameters under the null is estimated to be at most the number of parameters in ***θ*** that lie in the interior of the null parameter space.

For (ii), to define weak identification, let ***θ*** be a vector containing the null parameters (the ***γ***’s, *β*’s, and *π*), then the Jacobian of the function ***θ*** → ***q*** is approximately low rank for some values of ***θ***, which results in issues for the asymptotic performance of the LRT [Kleibergen, 2005, Han and McCloskey, 2019]. To address this weak identifiability problem, we calculate the Jacobian matrix for the internal parameters at the MLE of the function ***θ*** → ***q***. The numeric rank of this Jacobian matrix is used as the number of free parameters under the null. By default, the numeric rank we used is the number of singular values that are at least one-thousandth as large as the largest singular value. This heuristic is based on the literature that shows that the number of parameters in an exactly non-identified parameter space is the exact rank of the Jacobian [Catchpole and Morgan, 1997, Viallefont et al., 1998, Schmittmann et al., 2010]. Thus, we use the approximate rank of the Jacobian to approximate the number of parameters in an weakly identified parameter space. This heuristic works well in practice (Section 3), though we have not seen its use in prior literature.

The above scenarios assume the parental genotypes are known. In the case that parental genotypes are not known, we first maximize the likelihood jointly over the parental genotypes and the unknown null parameters (the ***γ***’s, *β*’s, and *π*). We then proceed with the LRT using the MLEs of the parent genotypes as if they were the true parent genotypes.

### 2.3 Implementation features

All methods described in this manuscript are implemented in the user-friendly segtest R package on GitHub (https://github.com/dcgerard/segtest). The package includes additional features that may be useful for applied researchers:

1. Bayesian Information Criterion (BIC): segtest computes the BIC [Schwarz, 1978] of the fitted null model using the estimated number of parameters under the null. Researchers can visualize BIC distributions across loci to assess model suitability, particularly when segregation patterns are unknown (e.g., to determine whether partial preferential pairing is present). The model with lower BIC values is generally preferred.
2. Outlier Detection: For each individual, segtest calculates the posterior probability of having an anomalous genotype at a given locus, allowing researchers to treat such genotypes as missing in downstream analyses. This follows a standard mixture model approach. For a *K*-ploid individual, let *q*_0_*, …, q_K_* be genotype likelihoods under the assumption of no outliers, computed via (1). The likelihoods for an individual as a non-outlier (*f*_1_) and as an outlier (*f*_0_) are defined as *f*_1_ = *q_k_* and 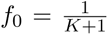 when the genotype *k* ∈ {0*, …, K*} is known. In the unknown genotype case, given individual genotype likelihoods *g*_0_*, …, g_K_*, these become 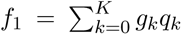 and 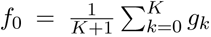. For a given outlier proportion *π*, the posterior probability that an individual is an outlier is then

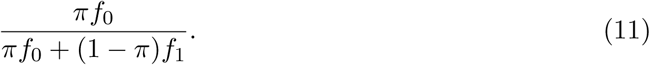

1. 3. Integration with updog: segtest includes helper functions to format output from updog [Gerard et al., 2018, Gerard and Ferrão, 2019] for direct input.
2. 4. Parallelization Support: The package supports parallel execution via the future package [Bengtsson, 2021], with additional compatibility for high-performance computing through future.batchtools.

## 3 Results

### 3.1 Null simulations

We assessed the type I error control of our new LRT and the polymapR test through simulations under the null hypothesis of no segregation distortion. The following parameters were varied:

- Ploidy: *K* ∈ {4, 6, 8}
- Parent genotypes: *ℓ*_1_ ∈ {0*, …, K*} and *ℓ*_2_ ∈ {*ℓ*_1_*, …, K*}
- Sample size: *n* ∈ {20, 200}
- Read depth: 10 (unknown genotype case) or infinity (known genotype case)
- Outlier proportion: *π* ∈ {0, 0.015, 0.03}
- Mixing proportions for the pairing configuration (for non-nulliplex and non-simplex loci):
  **–** If the number of mixture components is 2: (*γ*_1_*, γ*_2_) ∈ {(1, 0), (0.5, 0.5), (0, 1)}
  **–** For *K* = 8 and *ℓ* = 4, where three mixture components exist:
  (*γ*_1_*, γ*_2_*, γ*_3_) ∈ {(1, 0, 0), (0, 1, 0), (0, 0, 1), (1, 1, 0)*/*2, (1, 0, 1)*/*2, (0, 1, 1)*/*2, (1, 1, 1)*/*3}
- Double reduction adjustment for simplex loci: *β* ∈ {0*, a/*2*, a*}, where *a* is the maximum value under the complete equational segregation (CES) model in Table S2.

This yielded a total of 7920 distinct simulation scenarios to evaluate the null performance of our methods. For each scenario, genotype frequencies ***q***_0_ were determined. In the known genotype case, genotype counts were drawn from a multinomial distribution (9). In the unknown genotype case, these simulated counts were further processed to generate individual sequencing read counts according to the model of Gerard et al. [2018], assuming no allele bias, a sequencing error rate of 0.01, and an overdispersion parameter of 0.01. Genotype likelihoods were then obtained using the method of Gerard et al. [2018]. Each replication involved fitting either our LRT or the polymapR test [Bourke et al., 2018]. A total of 200 replications were performed per simulation scenario.

Figure 1 presents histograms of the type I error rates across the 7920 simulation scenarios at a nominal significance level of 0.05. Since the expected type I error rate should not exceed 0.05, any observed values above this threshold should only be attributable to the finite number of simulations. Specifically, the distribution of type I errors should be stochastically bounded by *X/*200, where *X* follows a binomial distribution with size 200 and success probability 0.05. For reference, Figure 1 includes the 99th percentile of this distribution as a blue dotted line. The results indicate that segtest adequately controls type I error for large sample sizes (*n* = 200) and for nearly all scenarios with small sample sizes (*n* = 20). In some small-sample scenarios, slight anticonservatism is observed, which is expected given that the LRT only asymptotically controls type I error.

**Figure 1:**
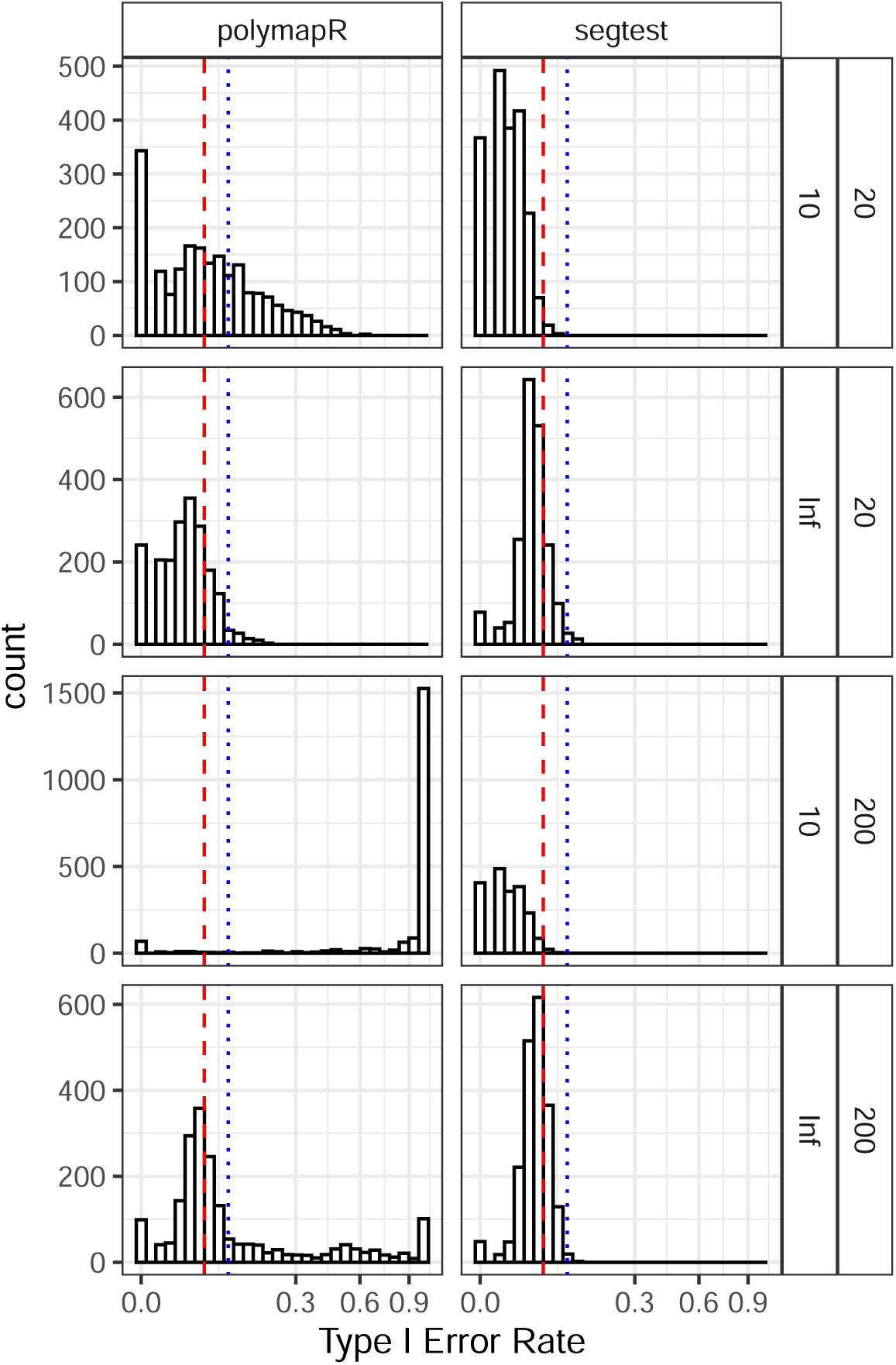
Histograms of type I error rates (on the square root scale) from the simulations in Section 3.1. Tests were conducted at a nominal significance level of 0.05. Results are stratified by sample size (*n* = 20 or 200) and read depth (10 or infinity) in the rows, and by method (segtest or polymapR) in the columns. Since the null hypothesis is true, the type I error rate should not exceed 0.05 (red dashed line), though small deviations are expected due to finite simulations. The blue dotted line marks the 99th percentile of expected variation under proper type I error control, based on binomial quantiles. Segtest controls type I error in all scenarios for large *n* = 200 and most for small *n* = 20, as expected given its asymptotic guarantees. In contrast, polymapR fails to control type I error at low read depths and for large sample sizes. It appears to control type I error at high read depths with small *n*, likely due to low power for small *n*.

In contrast, polymapR fails to control type I error in many scenarios, particularly for large sample sizes when genotypes are unknown. To illustrate this issue, Figure 2 presents results from select simulation scenarios. These simulations were conducted with a sample size of *n* = 200 octoploid (*K* = 8) offspring, where one parent had a genotype of 0 and the other had a genotype of 6. The figure contains quantile-quantile (Q-Q) plots of *p*-values against the uniform distribution, stratified by method (segtest or polymapR), read depth (10 or infinity), and pairing configuration mixing proportions for the second parent ((0.5, 0.5) or (1, 0)). Under the null hypothesis, the Q-Q plots should lie entirely at or above the *y* = *x* line (black), indicating appropriate type I error control.

**Figure 2:**
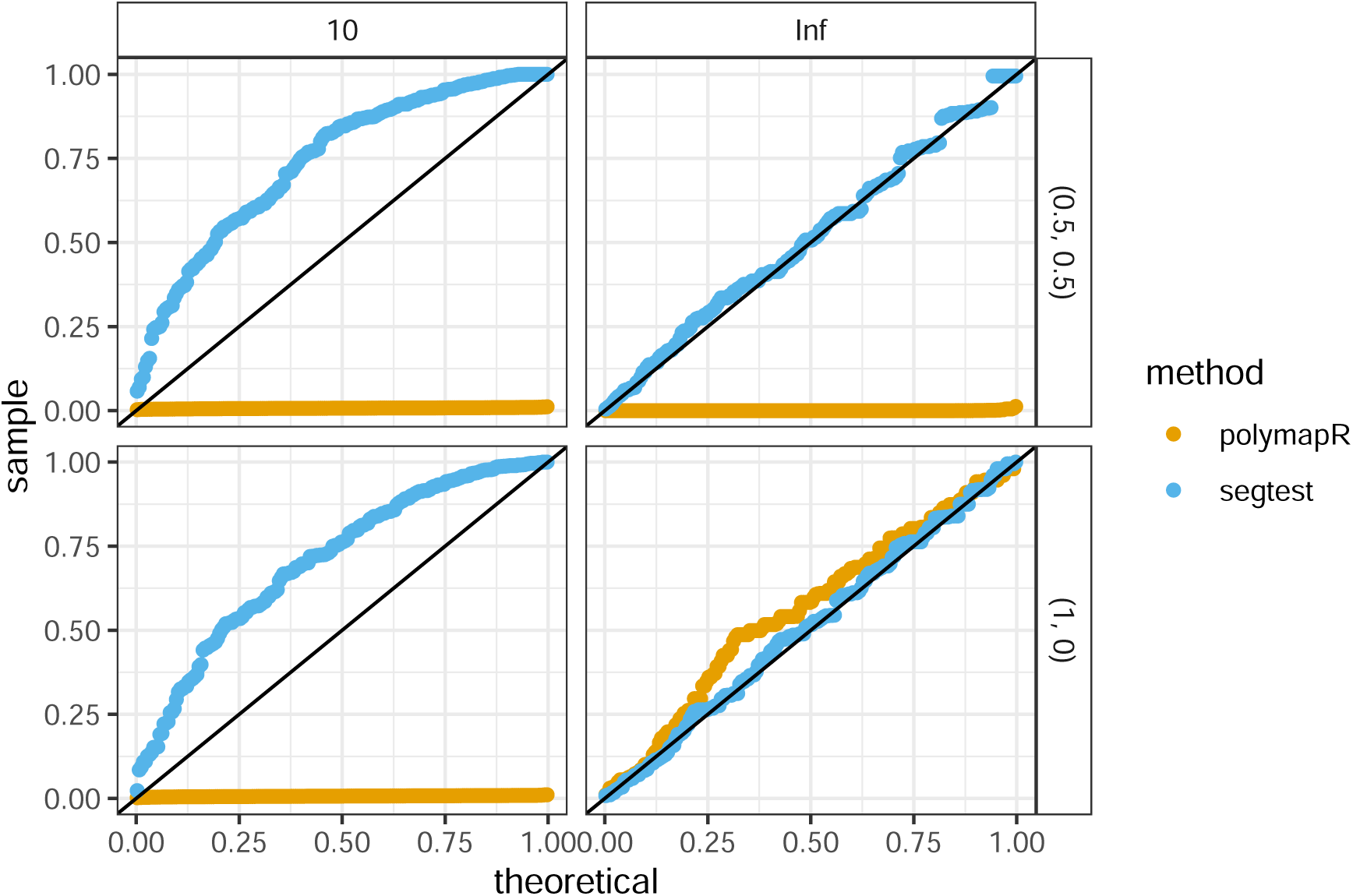
Example results from the null simulations in Section 3.1. Q-Q plots compare *p*-values from segtest (blue) and polymapR (orange) against the uniform distribution. Plots are stratified by read depth (columns) and pairing configuration (***γ***_2_, rows). This scenario assumes a ploidy of *K* = 8, parent genotypes *ℓ*_1_ = 0 and *ℓ*_2_ = 6, and a sample size of *n* = 200. Under the null, tests that properly control type I error should lie at or above the *y* = *x* line (black). PolymapR satisfies this condition only when ***γ***_2_ = (1, 0) (bottom row) and genotypes are fully known (infinite read depth). However, it fails to control type I error when preferential pairing is not absolute (top row) or when genotypes are uncertain (left column), even under its assumed null scenario. This is due to its ad hoc genotype uncertainty adjustment. In contrast, segtest consistently controls type I error across all scenarios.

Segtest satisfies this criterion in all scenarios, demonstrating proper type I error control at any significance level. In contrast, polymapR fails to control type I error in three cases. The method does not account for non-absolute preferential pairing, meaning that ***γ*** = (0.5, 0.5) represents an alternative scenario under its test, explaining its failure to control type I error at any read depth in this case. Additionally, polymapR fails to control type I error at low read depths even when absolute preferential pairing (***γ*** = (1, 0)) is assumed, a scenario that should fall within its null model. This failure arises because polymapR first estimates genotype counts before applying its known-genotype test, and this estimation process is biased [Gerard et al., 2025].

### 3.2 Robustness to double reduction

Since our method does not explicitly account for double reduction at non-simplex loci, we conducted simulations under the autopolyploid model of Huang et al. [2019], which allows for arbitrary levels of double reduction, though no preferential pairing. We varied the following parameters:

- Ploidy: *K* ∈ {4, 6, 8}
- Parent genotypes: *ℓ*_1_ ∈ {0*, …, K*} and *ℓ*_2_ ∈ {2*, …, K* − 2}
- Sample size: *n* ∈ {20, 200}
- Double reduction rate: ***α*** ∈ {**0**, ***α****_m_/*2, ***α****_m_*}, where ***α****_m_* is the maximum double reduction rate(s) under the CES model [Huang et al., 2019]
- Read depth: 10 (unknown genotype case) or infinity (known genotype case)

The outlier proportion was fixed at 0 (no outliers), resulting in 852 distinct simulation scenarios. Data were simulated as described in Section 3.1. For each replication, we applied the LRT assuming no outliers, the LRT allowing for outliers, and the polymapR test [Bourke et al., 2018], conducting 200 replications per scenario.

Figure 3 presents histograms of type I error rates at a nominal level of 0.05 across the simulated scenarios. When double reduction was absent, segtest, regardless of whether it accounted for outliers, maintained appropriate type I error control. In contrast, polymapR failed to control type I error at low read depths, consistent with the findings in Section 3.1.

**Figure 3:**
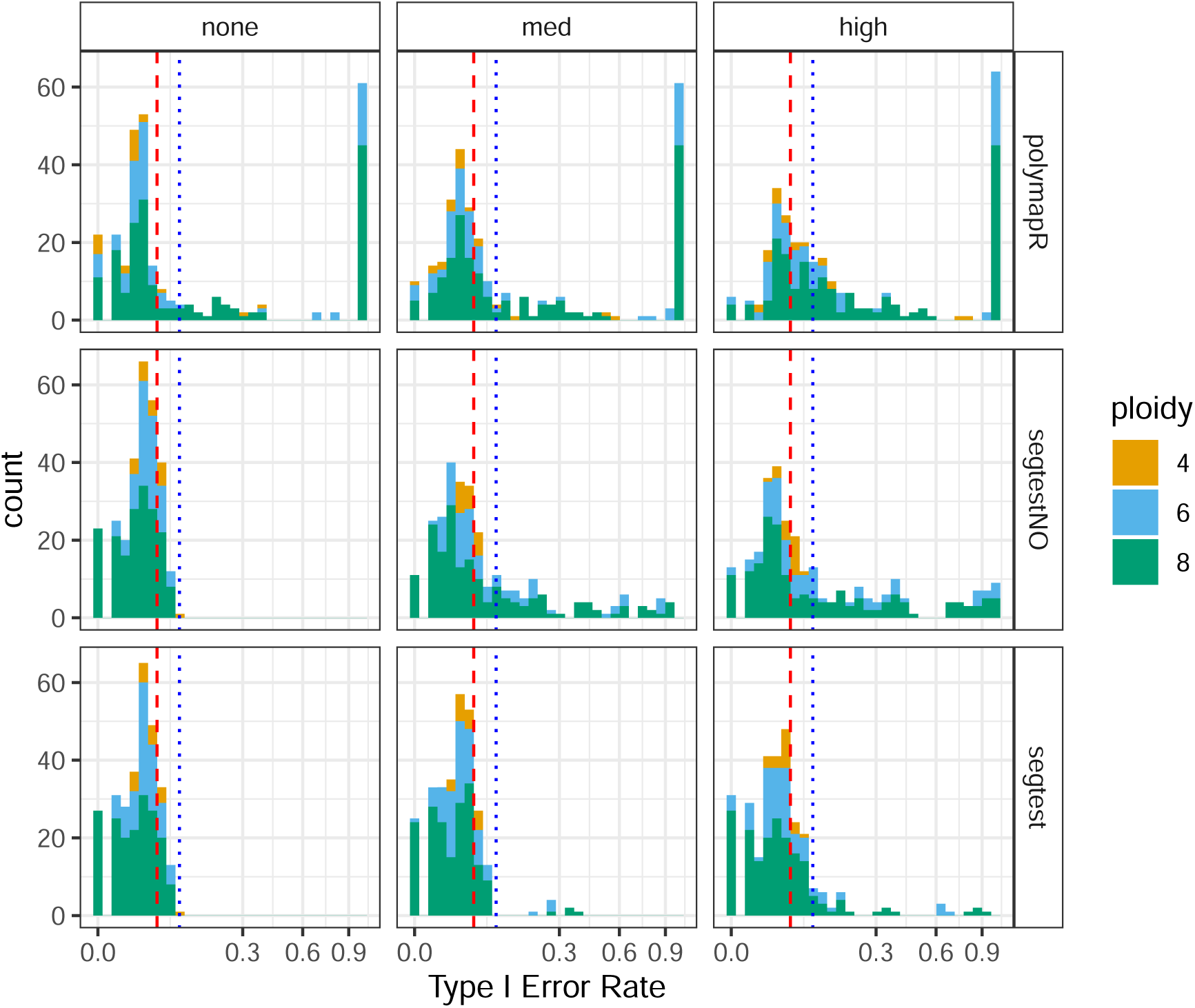
Robustness to double reduction from Section 3.2. Histograms of Type I error rates (square root scale) at a nominal significance level of 0.05 (red dashed line). The blue dotted line indicates the 99th percentile of expected variation under proper Type I error control, based on binomial quantiles. Results are stratified by the level of double reduction (none, medium, or high) along the columns and by method (polymapR, segtest assuming no outliers, and segtest allowing for outliers) along the rows. Ploidy is color-coded. Among the tested methods, only segtest with outlier accommodation remains relatively robust to the effects of double reduction, particularly at moderate levels.

At moderate levels of double reduction, only segtest with outlier accommodation remained relatively robust. There were eight simulation scenarios where segtest exhibited poor type I error control (above the 99th percentile of the Binomial(200, 0.05) distribution). These cases exclusively involved nulliplex-by-duplex parental genotypes at ploidies 6 and 8, with large sample sizes (*n* = 200) and known genotypes (Table S3). Notably, genotype uncertainty appeared to render the test conservative enough (see Section 3.1) to maintain adequate type I error control in the presence of moderate double reduction. Genotype uncertainty is the most common scenario in applied work [Gerard et al., 2018, Gerard and Ferrão, 2019].

At extreme levels of double reduction, only segtest with outlier handling performed reasonably well, though 30 of the 852 scenarios still exhibited poor type I error control. In such cases, type I error inflation was more pronounced. Though, extremely high double reduction rates are believed to be rare [Stift et al., 2008, Bomblies et al., 2016]. And in practice, such deviations would likely be identified by researchers in applied settings.

### 3.3 Power analysis

In this section, we evaluate the power of our methods under segregation distortion. We simulated genotype frequencies for ploidies *K* ∈ {4, 6, 8} under two alternative scenarios:

1. Easy scenario: The genotype frequency vector ***q*** was drawn uniformly from the *K*-dimensional unit simplex.
2. Hard scenario: Gamete frequency vectors ***p***_1_ and ***p***_2_ were first drawn independently and uniformly from the *K/*2-dimensional unit simplex and then convolved (1) to obtain ***q***.

The hard scenario generates alternative cases that more closely resemble the null, as the genotype frequencies are obtained by convolving randomly generated gamete frequencies, mirroring the null model structure. We further varied sample size (*n* ∈ {20, 200}) and read depth (10 or infinite). For each of 1000 replications, we simulated ***q***, generated data as described in Section 3.1, and applied both the LRT from segtest and the polymapR test.

Figure 4 presents the power of segtest and polymapR at a nominal significance level of 0.05. Because segtest properly controls type I error while polymapR does not, some reduction in power for segtest is expected. However, we find that segtest typically exhibits only a modest decrease in power, particularly for large samples (*n* = 200), where both methods perform well. In some hard scenarios with small *n*, segtest shows a more pronounced power reduction. Nonetheless, hypothesis testing requires valid type I error control, which segtest ensures, whereas polymapR does not. For sufficiently large sample sizes, segtest remains consistent and maintains power against all tested scenarios.

**Figure 4:**
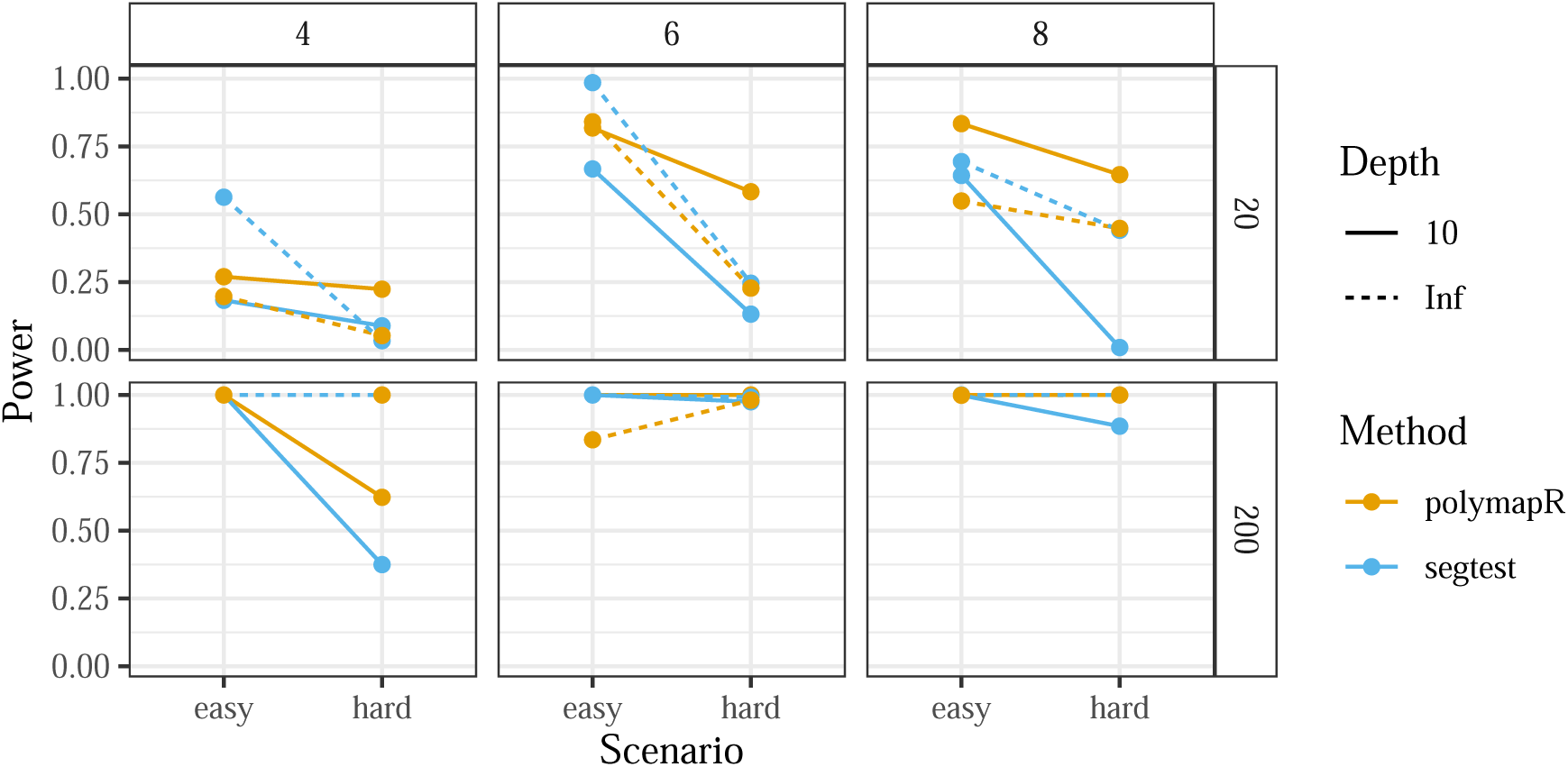
Results of the alternative simulations from Section 3.3. The figure plots the power (*y*-axis) of segtest and polymapR (color) under scenarios where segregation distortion is present. Tests were performed at a nominal significance level of 0.05 across varying sample sizes (*n* ∈ {20, 200}) (row facets), ploidies (*K* ∈ {4, 6, 8}) (column facets), read depths (10 or infinite) (line type), and alternative scenarios (easy or hard) (*x*-axis). segtest exhibits strong power for large *n* and maintains reasonable power in most small *n* scenarios. While segtest has slightly lower power than polymapR in most settings, this reduction is expected due to its ability to control type I error, which polymapR does not.

### 3.4 A hexaploid F1 population

To validate our methods, we analyzed the hexaploid sweetpotato (*Ipomoea batatas*, 2n = 6x = 90) mapping population from Mollinari et al. [2020], which consists of next-generation sequencing data from 315 full-sib individuals derived from a cross between the Beauregard and Tanzania varieties. These data include all SNPs from chromosome 8 (52,125 SNPs). After filtering out multiallelic loci (51,988 SNPs remaining), we performed genotyping using the method of Gerard et al. [2018] and Gerard and Ferrão [2019] with the “f1” model.

We tested for segregation distortion in these data using three different procedures. The first method emulated the default behavior of mappoly [Mollinari et al., 2020], following this procedure:

1. Filtered out SNPs estimated to be nulliplex in both parents (46,434 SNPs remaining).
2. For each SNP, filtered out individuals with a maximum genotype posterior probability below

0.95 (the default behavior of mappoly).

1. 3. Filtered out SNPs with an average read depth less than 20 (16,976 SNPs remaining), as done in Mollinari et al. [2020].
2. 4. Filtered out SNPs with at least 25% missing individuals (3,448 SNPs remaining), as done in Mollinari et al. [2020].

For the remaining SNPs, we used the posterior mode genotype as the “known” genotype and ran chi-squared tests for segregation distortion, assuming polysomic inheritance with bivalent pairing.

The second method used the checkF1() function from polymapR [Bourke et al., 2018]. Because this function accounts for genotype uncertainty, we only applied two filters: filtering out SNPs estimated to be nulliplex in both parents (46,434 SNPs remaining) and SNPs with an average read depth below 20 (16,105 SNPs remaining). We then applied the polymapR test using genotype posterior probabilities.

The third method applied our new approach based on genotype likelihoods, implemented in the segtest software. We applied the same pre-filters as in polymapR, analyzing 16,105 SNPs. We ran our new likelihood ratio test (Section 2.2) using three null models: our general model (Section 2.1), a polysomic inheritance model that allows for double reduction [Huang et al., 2019], and a polysomic inheritance model without double reduction [Serang et al., 2012]. For each locus, we computed the BIC of the null model (Section 2.3). We compared BICs pairwise at loci where the more complex null model had a *p*-value greater than 0.1. The polysomic model had a lower BIC than the general model 59.74% of the time (95% CI: 58.70% to 60.77%), the double reduction model had a lower BIC than the general model 55.85% of the time (95% CI: 54.80% to 56.90%), and the double reduction model had a lower BIC than the simpler polysomic model 51.66% of the time (95% CI: 50.57% to 52.75%). These results suggest that the polysomic inheritance model allowing for double reduction best fits these data, and thus tests from segtest based on this model are used in the comparisons that follow.

Histograms of the *p*-values from the three methods are shown in Figure 5. The *p*-value density for polymapR exhibits a mode at 1, which is generally not expected for unbiased *p*-values [Storey and Tibshirani, 2003]. In contrast, both mappoly and segtest display the more typical monotonically decreasing *p*-value densities. When controlling the false discovery rate at level 0.05, mappoly identifies far fewer SNPs (2,859 SNPs) not in segregation distortion than polymapR (13,796 SNPs) or segtest (10,058 SNPs). This suggests that mappoly’s filtering procedure, which does not account for genotype uncertainty, may be overly conservative, potentially discarding informative SNPs and leading to unnecessary data loss. Among SNPs in common after pre-filtering, the *p*-values from segtest and mappoly are correlated at 0.6996, those from segtest and polymapR at 0.4311, and those from polymapR and mappoly at 0.8169.

**Figure 5:**
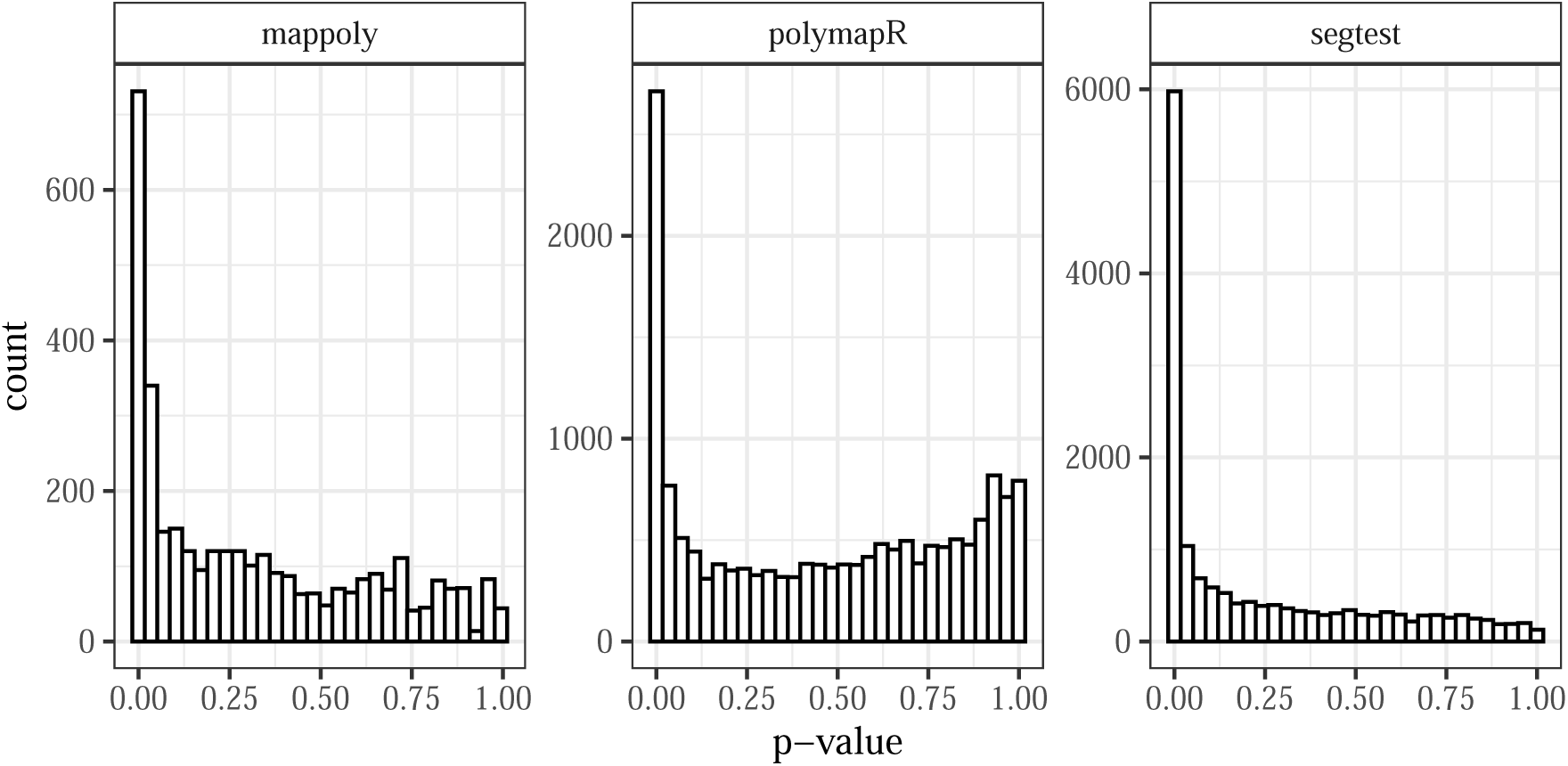
Histograms of the *p*-values from the segregation distortion tests performed by mappoly, polymapR, and segtest on the sweetpotato data described in Section 3.4. Note that the *y*-axes differ between facets.

It is informative to examine cases where the methods disagree. Figure S1 shows genotype plots [Gerard et al., 2018] for four SNPs where mappoly produces *p*-values less than 0.05 while segtest produces *p*-values greater than 0.1, and four SNPs with the opposite pattern. The *p*-values for these SNPs are listed in Table S4. Because mappoly filters out individuals with low genotype posterior probabilities before testing for segregation distortion, this alters the observed genotype frequencies and may lead to anticonservative results. For example, if we include those individuals in the mappoly test, the *p*-value becomes non-significant (Table S4). Conversely, segtest often yields small *p*-values for SNPs with many outlier genotypes, which mappoly has pre-filtered as “invalid.” These outlier individuals typically have low (or no) reference reads and may reflect null alleles [Gautier et al., 2013]. When we remove individuals with fewer than 30 reference reads, segtest produces larger *p*-values (Table S4). Whether such SNPs should be flagged is debatable, but in cases like SNP S8 2212186, where 21 of 315 individuals (≈6.7%) have “invalid” genotypes, flagging seems appropriate. Alternatively, if a researcher does not wish to flag such SNPs, instead of mixing the genotype frequencies with a discrete uniform in (8), we can mix them with an outlier distribution that one would expect in the presence of null alleles (a pointmass at a genotype of 0). When doing so, we again obtain much larger *p*-values (Table S4).

We conducted a similar comparison between polymapR and segtest. Figure S2 displays four SNPs where polymapR indicates segregation distortion (*p*-value *<* 0.05) but segtest does not (*p*-value *>* 0.1), and four SNPs where the reverse is true. The *p*-values from both methods are reported in Table S5. Like mappoly, polymapR removes “invalid” genotypes prior to testing. Thus, we again find that outliers explain the SNPs flagged by segtest but not by polymapR. When individuals with low reference reads are excluded, segtest returns much larger *p*-values (Table S5). Segtest also produces much larger *p*-values when the outlier distribution is a pointmass at a genotype of 0, thereby accounting for null alleles (Table S5). To understand cases where polymapR produces low *p*-values but segtest does not, we refer to Table S6, which shows the null and alternative genotype frequencies compared by each method. Both use the same alternative model, so the alternative frequencies are similar, differing only slightly due to polymapR’s ad hoc estimation versus segtest’s maximum likelihood approach [Li, 2011]. The key difference lies in the null frequencies. Segtest can model double reduction, allowing its null frequencies to better match the alternative frequencies than those used by polymapR. In these cases, the ability to account for double reduction appears to explain why segtest does not detect segregation distortion, while polymapR does.

## 4 Discussion

In this study, we developed statistical tests for segregation distortion in F1 populations of arbitrary (even) ploidy. Our methods account for varying levels of partial preferential pairing, double reduction at simplex loci, and a proportion of outliers. Effective modeling of outliers enhances robustness to moderate levels of double reduction at non-simplex loci. Additionally, our approach accommodates genotype uncertainty through genotype likelihoods. While our methods have only asymptotic guarantees for controlling type I error, we improve finite-sample performance by adaptively estimating the degrees of freedom, by counting boundary parameters and approximating the rank of a particular Jacobian. As a result, our LRT controls type I error reasonably well for sample sizes as small as 20 and performs robustly for samples of size 200.

For tetraploids, our methods fully and jointly account for double reduction and (partial) preferential pairing, as they generalize the approach of Gerard et al. [2025]. That is, due to the unidentifiability of the double reduction rate and the preferential pairing parameters for tetraploids, as described in Gerard et al. [2025], one can without loss of generality set the double reduction rate to 0 (at duplex loci), which is essentially what our method does for tetraploids in this manuscript. However, for ploidies greater than four, our model does not explicitly account for double reduction at non-simplex loci. For example, in hexaploids (*K* = 6), double reduction could theoretically produce gamete genotypes of three at a duplex locus, but our model does not allow for this possibility (Section S2). Nonetheless, by incorporating a small proportion of outliers, our approach demonstrates robustness to moderate levels of double reduction (Section 3.2). Increasing the maximum outlier proportion may further enhance type I error control in cases of extreme double reduction, though this would likely come at the cost of reduced power.

Our results here focus on scenarios that permit arbitrary levels of preferential pairing while explicitly modeling double reduction at simplex loci and allowing for a small fraction of outliers. However, our segtest package offers greater flexibility, enabling segregation distortion testing under additional assumptions:

1. True autopolyploidy, assuming either complete bivalent pairing (no double reduction) or arbitrary levels of double reduction, modeled as in Huang et al. [2019].
2. True allopolyploidy with complete preferential pairing.
3. An unknown ploidy structure, where the data could correspond to either a true allopolyploid or a bivalent-pairing autopolyploid. This is the assumption used by polymapR.

Additionally, segtest allows for testing without permitting outliers, modeling segregation distortion when any individual exhibits an “impossible” genotype. Each of these scenarios represents a special case of the general model studied here. If researchers have prior knowledge about the expected segregation patterns in their organism, they should specify the most appropriate null model. The segtest package also provides the BIC [Schwarz, 1978] for a given model (Section 2.3), which can aid researchers in determining the most suitable segregation type for their data.

We sought to improve the finite sample performance of the likelihood ratio test by adaptively estimating its degrees of freedom. An alternative approach would have been to use the bootstrap [Efron, 1979], which is known to provide highly accurate finite-sample results across many model classes [Efron, 1987]. However, we did not pursue this approach primarily due to computational constraints. Our tests are designed for genomic applications, where they may be applied thousands or even millions of times within a single study. Any method requiring more than approximately one second per test imposes an undue burden on applied researchers. Nonetheless, as computational power continues to increase, the bootstrap remains a promising avenue for future exploration.

Mappoly assumes genotypes are known when performing genetic mapping, and so it is natural that it filters out individuals with high genotype uncertainty. However, our results in Section 3.4 suggest that filtering individuals prior to testing for segregation distortion can lead to biased tests and unnecessary data loss. We recommend reversing the order of operations: rather than filtering out uncertain genotypes before testing, researchers should first apply a test that accounts for genotype uncertainty (e.g., our approach), and then filter out individuals with high uncertainty. Understanding how different data quality control steps influence downstream applications such as genetic mapping is an important question for future study.

## Acknowledgments

Most analyses were performed using the R statistical language [R Core Team, 2025].

## Data accessibility

All methods described in this paper are implemented in the segtest R package on the GitHub:

https://github.com/dcgerard/segtest

All analysis scripts to reproduce the results of this paper are available on GitHub:

https://github.com/dcgerard/seganal

https://github.com/dcgerard/seganal_data

The data used in this manuscript is from Mollinari et al. [2020] and is available on Figshare:

https://doi.org/10.25387/g3.10255844

## Supplementary material

Additional figures, tables, and theoretical details are available in the Supplementary Material online.

## Author contributions

DG developed the methodology, wrote the software, analyzed the data, implemented the study, and wrote the manuscript.

GBA analyzed the data, implemented the study, and wrote the manuscript. GDSP implemented the study and wrote the manuscript.

AAFG implemented the study and wrote the manuscript.

## Notes

https://github.com/dcgerard/segtest

https://github.com/dcgerard/seganal

https://github.com/dcgerard/seganal_data

### Competing Interest Statement

The authors have declared no competing interest.

