## Supplementary Materials for "Tests for segregation distortion in higher ploidy F1 populations"

**Abstract**

This document contains additional theoretical considerations, derivations, and figures to supplement the manuscript "Tests for segregation distortion in higher ploidy F1 populations".

### S1 Proof of theorem 1

*Proof.* For $p_0$, the minor allele is not in the gamete if it is not one of the $i$ double reduced alleles (probability $\binom{K-1}{i}/\binom{K}{i}$) and it is not one of the non-double-reduced alleles (probability $\binom{K-1-i}{K/2-2i}/\binom{K-i}{K/2-2i}$). We marginalize over the number of double reduced pairs to obtain

$$
\begin{aligned}
p_0 &= \sum_{i=0}^{\lfloor K/4 \rfloor} \alpha_i \frac{\binom{K-1}{i}\binom{K-1-i}{K/2-2i}}{\binom{K}{i}\binom{K-i}{K/2-2i}} && \text{(S1)}\\
&= \sum_{i=0}^{\lfloor K/4 \rfloor} \alpha_i \frac{(K-1)!}{i!(K-1-i)!}\frac{(K-1-i)!}{(K/2-2i)!(K/2+i-1)!}\frac{i!(K-i)!}{K!}\frac{(K/2-2i)!(K/2+i)!}{(K-i)!} && \text{(S2)}\\
&= \sum_{i=0}^{\lfloor K/4 \rfloor} \alpha_i \frac{K/2+i}{K} && \text{(S3)}\\
&= \frac{1}{2}\alpha_0 + \sum_{i=1}^{\lfloor K/4 \rfloor} \alpha_i \frac{K/2+i}{K} && \text{(S4)}\\
&= \frac{1}{2}\left(1 - \sum_{i=1}^{\lfloor K/4 \rfloor} \alpha_i\right) + \sum_{i=1}^{\lfloor K/4 \rfloor} \alpha_i \frac{K/2+i}{K} && \text{(S5)}\\
&= \frac{1}{2} + \sum_{i=1}^{\lfloor K/4 \rfloor} \alpha_i \left(\frac{K/2+i}{K} - \frac{1}{2}\right) && \text{(S6)}\\
&= \frac{1}{2} + \sum_{i=1}^{\lfloor K/4 \rfloor} \alpha_i \frac{i}{K}. && \text{(S7)}
\end{aligned}
$$

Similarly, for $p_1$, the minor allele shows up once in the gamete if it is not one of the $i$ double reduced alleles (probability $\binom{K-1}{i}/\binom{K}{i}$) and it *is* one of the non-double-reduced alleles (probability $\binom{K-1-i}{K/2-2i-1}/\binom{K-i}{K/2-2i}$). We marginalize over the number of double reduced pairs to obtain

$$
\begin{aligned}
p_1 &= \sum_{i=0}^{\lfloor K/4 \rfloor} \alpha_i \frac{\binom{K-1}{i}\binom{K-1-i}{K/2-2i-1}}{\binom{K}{i}\binom{K-i}{K/2-2i}} && \text{(S8)} \\
&= \sum_{i=0}^{\lfloor K/4 \rfloor} \alpha_i \frac{(K-1)!}{i!(K-1-i)!} \frac{(K-1-i)!}{(K/2-2i-1)!(K/2+i)!} \frac{i!(K-i)!}{K!} \frac{(K/2-2i)!(K/2+i)!}{(K-i)!} && \text{(S9)} \\
&= \sum_{i=0}^{\lfloor K/4 \rfloor} \alpha_i \frac{K/2-2i}{K} && \text{(S10)} \\
&= \frac{1}{2}\alpha_0 + \sum_{i=1}^{\lfloor K/4 \rfloor} \alpha_i \frac{K/2-2i}{K} && \text{(S11)} \\
&= \frac{1}{2}\left(1 - \sum_{i=1}^{\lfloor K/4 \rfloor} \alpha_i\right) + \sum_{i=1}^{\lfloor K/4 \rfloor} \alpha_i \frac{K/2-2i}{K} && \text{(S12)} \\
&= \frac{1}{2} + \sum_{i=1}^{\lfloor K/4 \rfloor} \alpha_i \left(\frac{K/2-2i}{K} - \frac{1}{2}\right) && \text{(S13)} \\
&= \frac{1}{2} - 2\sum_{i=1}^{\lfloor K/4 \rfloor} \alpha_i \frac{i}{K} && \text{(S14)}
\end{aligned}
$$

For $p_2$, we could derive its value from $p_0 + p_1 + p_2 = 1$. Or we can derive it from first principles. The minor allele shows up twice in the gamete if it is one of the $i$ double reduced alleles (probability $\binom{K-1}{i-1}/\binom{K}{i}$). Marginalizing over the number of double reduced pairs, we have

$$
\begin{aligned}
p_2 &= \sum_{i=1}^{\lfloor K/4 \rfloor} \alpha_i \frac{\binom{K-1}{i-1}}{\binom{K}{i}} && \text{(S15)} \\
&= \sum_{i=1}^{\lfloor K/4 \rfloor} \alpha_i \frac{(K-1)!}{(i-1)!(K-i)!} \frac{i!(K-i)!}{K!} && \text{(S16)} \\
&= \sum_{i=1}^{\lfloor K/4 \rfloor} \alpha_i \frac{i}{K}. && \text{(S17)}
\end{aligned}
$$

□

### S2 Our model for the gamete frequencies for ploidies 2, 4, 6, 8, 10, and 12

Given a parent dosage, $\ell$, below we list the model for the gamete frequencies $\boldsymbol{p} = (p_0, \ldots, p_{K/2})$ at a given ploidy $K$. Note that $\sum \gamma_i = 1$.

Ploidy = 2

- $\ell = 0$: $(1, 0)$
- $\ell = 1$: $(1, 1)/2$
- $\ell = 2$: $(0, 1)$

Ploidy = 4

- $\ell = 0$: $(1, 0, 0)$
- $\ell = 1$: $(\frac{1}{2} + \beta, \frac{1}{2} - 2\beta, \beta)$
- $\ell = 2$: $\gamma_1 \frac{1}{4}(1, 2, 1) + \gamma_2(0, 1, 0)$
- $\ell = 3$: $(\beta, \frac{1}{2} - 2\beta, \frac{1}{2} + \beta)$
- $\ell = 4$: $(0, 0, 1)$

Ploidy = 6

- $\ell = 0$: $(1, 0, 0, 0)$
- $\ell = 1$: $(\frac{1}{2} + \beta, \frac{1}{2} - 2\beta, \beta, 0)$
- $\ell = 2$: $\gamma_1 \frac{1}{4}(1, 2, 1, 0) + \gamma_2(0, 1, 0, 0)$
- $\ell = 3$: $\gamma_1 \frac{1}{8}(1, 3, 3, 1) + \gamma_2 \frac{1}{2}(0, 1, 1, 0)$
- $\ell = 4$: $\gamma_1 \frac{1}{4}(0, 1, 2, 1) + \gamma_2(0, 0, 1, 0)$
- $\ell = 5$: $(0, \beta, \frac{1}{2} - 2\beta, \frac{1}{2} + \beta)$
- $\ell = 6$: $(0, 0, 0, 1)$

Ploidy = 8

- $\ell = 0$: $(1, 0, 0, 0, 0)$
- $\ell = 1$: $(\frac{1}{2} + \beta, \frac{1}{2} - 2\beta, \beta, 0, 0)$
- $\ell = 2$: $\gamma_1 \frac{1}{4}(1, 2, 1, 0, 0) + \gamma_2(0, 1, 0, 0, 0)$
- $\ell = 3$: $\gamma_1 \frac{1}{8}(1, 3, 3, 1, 0) + \gamma_2 \frac{1}{2}(0, 1, 1, 0, 0)$
- $\ell = 4$: $\gamma_1 \frac{1}{16}(1, 4, 6, 4, 1) + \gamma_2 \frac{1}{4}(0, 1, 2, 1, 0) + \gamma_3(0, 0, 1, 0, 0)$
- $\ell = 5$: $\gamma_1 \frac{1}{8}(0, 1, 3, 3, 1) + \gamma_2 \frac{1}{2}(0, 0, 1, 1, 0)$
- $\ell = 6$: $\gamma_1 \frac{1}{4}(0, 0, 1, 2, 1) + \gamma_2(0, 0, 0, 1, 0)$
- $\ell = 7$: $(0, 0, \beta, \frac{1}{2} - 2\beta, \frac{1}{2} + \beta)$
- $\ell = 8$: $(0, 0, 0, 0, 1)$

Ploidy = 10

- $\ell = 0$: $(1, 0, 0, 0, 0, 0)$
- $\ell = 1$: $(\frac{1}{2} + \beta, \frac{1}{2} - 2\beta, \beta, 0, 0, 0)$
- $\ell = 2$: $\gamma_1 \frac{1}{4}(1, 2, 1, 0, 0, 0) + \gamma_2(0, 1, 0, 0, 0, 0)$

- $\ell = 3$: $\gamma_1 \frac{1}{8}(1,3,3,1,0,0) + \gamma_2 \frac{1}{2}(0,1,1,0,0,0)$
- $\ell = 4$: $\gamma_1 \frac{1}{16}(1,4,6,4,1,0) + \gamma_2 \frac{1}{4}(0,1,2,1,0,0) + \gamma_3(0,0,1,0,0,0)$
- $\ell = 5$: $\gamma_1 \frac{1}{32}(1,5,10,10,5,1) + \gamma_2 \frac{1}{8}(0,1,3,3,1,0) + \gamma_3 \frac{1}{2}(0,0,1,1,0,0)$
- $\ell = 6$: $\gamma_1 \frac{1}{16}(0,1,4,6,4,1) + \gamma_2 \frac{1}{4}(0,0,1,2,1,0) + \gamma_3(0,0,0,1,0,0)$
- $\ell = 7$: $\gamma_1 \frac{1}{8}(0,0,1,3,3,1) + \gamma_2 \frac{1}{2}(0,0,0,1,1,0)$
- $\ell = 8$: $\gamma_1 \frac{1}{4}(0,0,0,1,2,1) + \gamma_2(0,0,0,0,1,0)$
- $\ell = 9$: $(0,0,0,\beta,\frac{1}{2}-2\beta,\frac{1}{2}+\beta)$
- $\ell = 10$: $(0,0,0,0,0,1)$

Ploidy = 12

- $\ell = 0$: $(1,0,0,0,0,0,0)$
- $\ell = 1$: $(\frac{1}{2}+\beta,\frac{1}{2}-2\beta,\beta,\beta,0,0,0,0)$
- $\ell = 2$: $\gamma_1 \frac{1}{4}(1,2,1,0,0,0,0) + \gamma_2(0,1,0,0,0,0,0)$
- $\ell = 3$: $\gamma_1 \frac{1}{8}(1,3,3,1,0,0,0) + \gamma_2 \frac{1}{2}(0,1,1,0,0,0,0)$
- $\ell = 4$: $\gamma_1 \frac{1}{16}(1,4,6,4,1,0,0) + \gamma_2 \frac{1}{4}(0,1,2,1,0,0,0) + \gamma_3(0,0,1,0,0,0,0)$
- $\ell = 5$: $\gamma_1 \frac{1}{32}(1,5,10,10,5,1,0) + \gamma_2 \frac{1}{8}(0,1,3,3,1,0,0) + \gamma_3 \frac{1}{2}(0,0,1,1,0,0,0)$
- $\ell = 6$: $\gamma_1 \frac{1}{64}(1,6,15,20,15,6,1)+\gamma_2 \frac{1}{16}(0,1,4,6,4,1,0)+\gamma_3 \frac{1}{4}(0,0,1,2,1,0,0)+\gamma_4(0,0,0,1,0,0,0)$
- $\ell = 7$: $\gamma_1 \frac{1}{32}(0,1,5,10,10,5,1) + \gamma_2 \frac{1}{8}(0,0,1,3,3,1,0) + \gamma_3 \frac{1}{2}(0,0,0,1,1,0,0)$
- $\ell = 8$: $\gamma_1 \frac{1}{16}(0,0,1,4,6,4,1) + \gamma_2 \frac{1}{4}(0,0,0,1,2,1,0) + \gamma_3(0,0,0,0,1,0,0)$
- $\ell = 9$: $\gamma_1 \frac{1}{8}(0,0,0,1,3,3,1) + \gamma_2 \frac{1}{2}(0,0,0,0,1,1,0)$
- $\ell = 10$: $\gamma_1 \frac{1}{4}(0,0,0,0,1,2,1) + \gamma_2(0,0,0,0,0,1,0)$
- $\ell = 11$: $(0,0,0,0,\beta,\frac{1}{2}-2\beta,\frac{1}{2}+\beta)$
- $\ell = 12$: $(0,0,0,0,0,0,1)$

### S3 Supplementary tables, figures, and procedures

| Parent Dosage | Ploidy | | | | | |
|---|---|---|---|---|---|---|
| | 2 | 4 | 6 | 8 | 10 | 12 |
| 0 | (1,0) | (1,0,0) | (1,0,0,0) | (1,0,0,0,0) | (1,0,0,0,0,0) | (1,0,0,0,0,0,0) |
| 1 | (1,1)/2 | (1,1,0)/2 | (1,1,0,0)/2 | (1,1,0,0,0)/2 | (1,1,0,0,0,0)/2 | (1,1,0,0,0,0,0)/2 |
| 2 | (0,1) | (1,2,1)/4<br>(0,1,0) | (1,2,1,0)/4<br>(0,1,0,0) | (1,2,1,0,0)/4<br>(0,1,0,0,0) | (1,2,1,0,0,0)/4<br>(0,1,0,0,0,0) | (1,2,1,0,0,0,0)/4<br>(0,1,0,0,0,0,0) |
| 3 | - | (0,1,1)/2 | (1,3,3,1)/8<br>(0,1,1,0)/2 | (1,3,3,1,0)/8<br>(0,1,1,0,0)/2 | (1,3,3,1,0,0)/8<br>(0,1,1,0,0,0)/2 | (1,3,3,1,0,0,0)/8<br>(0,1,1,0,0,0,0)/2 |
| 4 | - | (0,0,1) | (0,1,2,1)/4<br>(0,0,1,0) | (1,4,6,4,1)/16<br>(0,1,2,1,0)/4<br>(0,0,1,0,0) | (1,4,6,4,1,0)/16<br>(0,1,2,1,0,0)/4<br>(0,0,1,0,0,0) | (1,4,6,4,1,0,0)/16<br>(0,1,2,1,0,0,0)/4<br>(0,0,1,0,0,0,0) |
| 5 | - | - | (0,0,1,1)/2 | (0,1,3,3,1)/8<br>(0,0,1,1,0)/2 | (1,5,10,10,5,1)/32<br>(0,1,3,3,1,0)/8<br>(0,0,1,1,0,0)/2 | (1,5,10,10,5,1,0)/32<br>(0,1,3,3,1,0,0)/8<br>(0,0,1,1,0,0,0)/2 |
| 6 | - | - | (0,0,0,1) | (0,0,1,2,1)/4<br>(0,0,0,1,0) | (0,1,4,6,4,1)/16<br>(0,0,1,2,1,0)/4<br>(0,0,0,1,0,0) | (1,6,15,20,15,6,1)/64<br>(0,1,4,6,4,1,0)/16<br>(0,0,1,2,1,0,0)/4<br>(0,0,0,1,0,0,0) |
| 7 | - | - | - | (0,0,0,1,1)/2 | (0,0,1,3,3,1)/8<br>(0,0,0,1,1,0)/2 | (0,1,5,10,10,5,1)/32<br>(0,0,1,3,3,1,0)/8<br>(0,0,0,1,1,0,0)/2 |
| 8 | - | - | - | (0,0,0,0,1) | (0,0,0,1,2,1)/4<br>(0,0,0,0,1,0) | (0,0,1,4,6,4,1)/16<br>(0,0,0,1,2,1,0)/4<br>(0,0,0,0,1,0,0) |
| 9 | - | - | - | - | (0,0,0,0,1,1)/2 | (0,0,0,1,3,3,1)/8<br>(0,0,0,0,1,1,0)/2 |
| 10 | - | - | - | - | (0,0,0,0,0,1) | (0,0,0,0,1,2,1)/4<br>(0,0,0,0,0,1,0) |
| 11 | - | - | - | - | - | (0,0,0,0,0,1,1)/2 |
| 12 | - | - | - | - | - | (0,0,0,0,0,0,1) |

Table S1: Gamete frequencies for true allopolyploids. Each cell contains the possible gamete frequencies $(p_0, p_1, \ldots, p_{K/2})$ of a true allopolyploid for a given ploidy $K$ (column) and a given parental dosage (row).

| Meiosis Model | Ploidy | | | | |
|---|---|---|---|---|---|
| | 4 | 6 | 8 | 10 | 12 |
| PRCS | $1/28 \approx 0.03571$ | $1/22 \approx 0.04545$ | $1/20 = 0.05$ | $1/19 \approx 0.05263$ | $5/92 \approx 0.05435$ |
| CES | $1/24 \approx 0.04167$ | $1/20 = 0.05$ | $3/56 \approx 0.05357$ | $1/18 \approx 0.05556$ | $5/88 \approx 0.05682$ |

Table S2: Upper bounds on $\beta$ from (7) under either the complete equational segregation model (CES) (where whole arms of sister chromatids are exchanged) [Mather, 1935] or the pure random chromatid segregation model (PRCS) (where chromatids act independently and segregate into gametes with equal probability) [Haldane, 1930].

| $n$ | Ploidy | Read Depth | $\ell_1$ | $\ell_2$ | Type I Error Rate |
|---|---|---|---|---|---|
| 200 | 8 | Inf | 0 | 2 | 0.38 |
| 200 | 8 | Inf | 8 | 2 | 0.35 |
| 200 | 8 | Inf | 0 | 6 | 0.34 |
| 200 | 8 | Inf | 8 | 6 | 0.28 |
| 200 | 6 | Inf | 0 | 2 | 0.26 |
| 200 | 6 | Inf | 0 | 4 | 0.26 |
| 200 | 6 | Inf | 6 | 2 | 0.26 |
| 200 | 6 | Inf | 6 | 4 | 0.20 |

Table S3: Eight simulation scenarios for `segtest` with poor type I error control under moderate double reduction (Section 3.2). All scenarios involve duplex-by-nullplex crosses, known genotypes, and large sample sizes.

| SNP | `mappoly` | `segtest` | `mappoly` All | `segtest` Less | `segtest` Null Alleles |
|---|---|---|---|---|---|
| S8_2301244 | 0.01 | 0.21 | 0.19 | | |
| S8_2301256 | 0.0034 | 0.11 | 0.12 | | |
| S8_5495374 | 0.0053 | 0.9 | 0.18 | | |
| S8_16038595 | 0.031 | 0.33 | 0.3 | | |
| S8_460771 | 0.12 | 8.9E-08 | | 0.19 | 0.053 |
| S8_2212186 | 0.13 | 2.8E-10 | | 0.18 | 0.00012 |
| S8_4866982 | 0.51 | 1.7E-07 | | 0.22 | 1 |
| S8_18370562 | 0.36 | 0.045 | | 0.3 | 0.62 |

Table S4: $P$-values for SNPs where the results from `mappoly` and `segtest` differ. The "`mappoly`" and "`segtest`" columns show $p$-values under each method's default behavior. The "`mappoly` All" column shows $p$-values from `mappoly` when including individuals that are normally filtered out based on maximum posterior probability. The "`segtest` Less" column shows $p$-values from `segtest` after removing individuals with low reference read counts. The "`segtest` Null Alleles" columns shows the $p$-values from `segtest` when using an outlier distribution that accounts for null alleles.

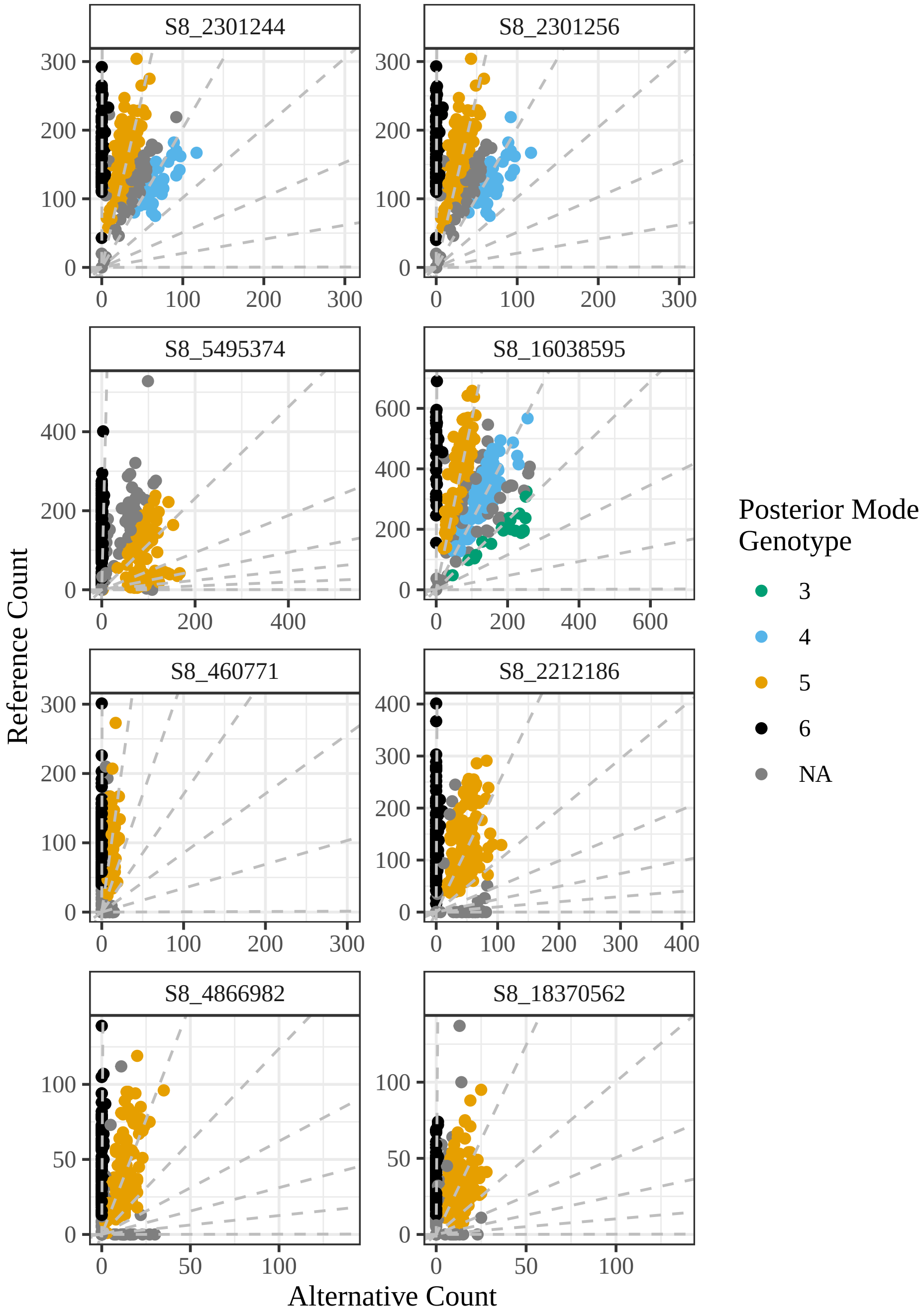


Figure S1: Genotype plots [Gerard et al., 2018] for a sample of SNPs where the results from `mappoly` and `segtest` differ. Alternative allele counts are on the $x$-axis, and reference allele counts are on the $y$-axis. Individuals are color-coded by posterior mode genotype. Missing values ("NA") indicate individuals that were filtered out by `mappoly`, though `segtest` includes all individuals in its test. The top four SNPs have low $p$-values from `mappoly` and high $p$-values from `segtest`. The bottom four SNPs show the opposite pattern, with low $p$-values from `segtest` and high $p$-values from `mappoly`.

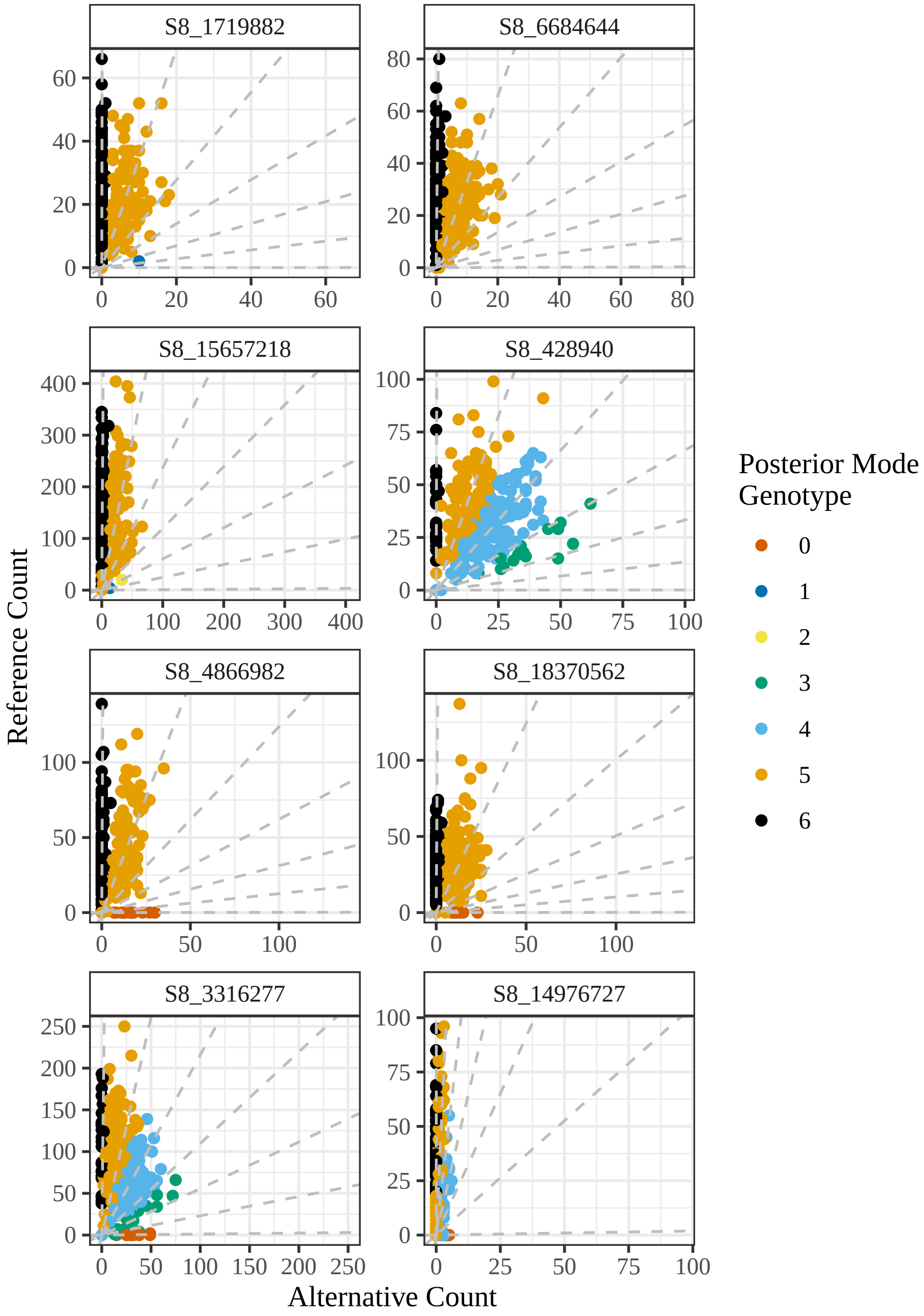


Figure S2: Genotype plots [Gerard et al., 2018] for a sample of SNPs where the results from `polymapR` and `segtest` differ. Alternative allele counts are on the $x$-axis, and reference allele counts are on the $y$-axis. Individuals are color-coded by posterior mode genotype. The top four SNPs have low $p$-values from `polymapR` and high $p$-values from `segtest`. The bottom four SNPs show the opposite pattern, with low $p$-values from `segtest` and high $p$-values from `polymapR`.

| SNP | `polymapR` | `segtest` | `segtest` Less | `segtest` Null Alleles |
|---|---:|---:|---:|---:|
| S8_1719882 | 0.0096 | 0.56 | | |
| S8_6684644 | 0.23 | 0.12 | | |
| S8_15657218 | 0.0073 | 0.55 | | |
| S8_428940 | 0.0012 | 0.8 | | |
| S8_4866982 | 0.41 | 1.7E-07 | 0.19 | 1 |
| S8_18370562 | 0.63 | 0.045 | 0.57 | 0.62 |
| S8_3316277 | 1 | 0.00046 | 0.35 | 0.37 |
| S8_14976727 | 0.93 | 1.4E-10 | 0.036 | 0.03 |

Table S5: $P$-values for SNPs where the results from `polymapR` and `segtest` differ. The "`polymapR`" and "`segtest`" columns show $p$-values under each method's default behavior. The "`segtest` Less" column shows $p$-values from `segtest` after removing individuals with low reference read counts. The "`segtest` Null Alleles" columns shows the $p$-values from `segtest` when using an outlier distribution that accounts for null alleles.

| | | Genotype | | | | | | |
|---|---|---:|---:|---:|---:|---:|---:|---:|
| SNP | Scenario | 0 | 1 | 2 | 3 | 4 | 5 | 6 |
| S8_15657218 | `polymapR` Alt | 0 | 0 | 0 | 0 | 0 | .425 | .575 |
| | `polymapR` Null | 0 | 0 | 0 | 0 | 0 | .500 | .500 |
| | `segtest` Alt | 0 | 0 | .006 | 0 | .036 | .376 | .582 |
| | `segtest` Null | .001 | .001 | .001 | .001 | .051 | .398 | .546 |
| S8_1719882 | `polymapR` Alt | 0 | 0 | 0 | 0 | 0 | .427 | .573 |
| | `polymapR` Null | 0 | 0 | 0 | 0 | 0 | .500 | .500 |
| | `segtest` Alt | 0 | 0 | .003 | 0 | .024 | .393 | .580 |
| | `segtest` Null | 0 | 0 | 0 | 0 | .047 | .407 | .545 |
| S8_428940 | `polymapR` Alt | 0 | 0 | 0 | .064 | .447 | .394 | .095 |
| | `polymapR` Null | 0 | 0 | 0 | .050 | .450 | .450 | .050 |
| | `segtest` Alt | 0 | 0 | .003 | .072 | .446 | .382 | .097 |
| | `segtest` Null | 0 | 0 | 0 | .091 | .409 | .409 | .091 |
| S8_6684644 | `polymapR` Alt | 0 | 0 | 0 | 0 | 0 | .424 | .576 |
| | `polymapR` Null | 0 | 0 | 0 | 0 | 0 | .500 | .500 |
| | `segtest` Alt | .003 | 0 | 0 | .024 | .001 | .392 | .581 |
| | `segtest` Null | .002 | .002 | .002 | .002 | .051 | .397 | .545 |

Table S6: Estimated genotype frequencies under different methods and hypotheses for four SNPs where `segtest` indicates no segregation distortion, but `polymapR` indicates strong segregation distortion. "`polymapR` Alt" contains `polymapR`'s estimated empirical genotype frequencies from genotype posterior probabilities [see Gerard et al., 2025, for a summary]. "`polymapR` Null" contains the expected genotype frequencies under polysomic inheritance with bivalent pairing. "`segtest` Alt" contains the genotype frequencies estimated using the method of Li [2011]. And "`segtest` Null" contains the estimated genotype frequencies under the null model from Section 2.1.
